# Low genetic variability and high isolation of a post-harvest South American pinniped population as revealed by genome-wide SNPs

**DOI:** 10.1101/2023.12.15.571663

**Authors:** Diego M. Peralta, Ezequiel A. Ibañez, Sergio Lucero, Humberto L. Cappozzo, Santiago G. Ceballos, Juan I. Túnez

**Author notes:** Corresponding autor: Diego M. Peralta,. Present address: Instituto de Ecología, UNAM. C.U., Coyoacán, 04510, CDMX, México. These authors contributed equally to this work (Santiago G. Ceballos, Juan I. Túnez).

## Abstract

*Otaria flavescens* has been one of the most heavily exploited pinnipeds during the last 200 years with depletions of about 90% in some colonies. After the prohibition on sealing in South America, populations became stabilized except for the Uruguayan population, which showed a constant decrease. The underlying causes of its decline have remained unknown. This study used the RAD-seq approach to assess the variability and connectivity of some of the most overexploited sea lion colonies in the Atlantic Ocean. Our results revealed low allelic richness, nucleotide diversity and heterozygosity in the Uruguayan population and evidence of complete isolation from the Argentinean populations under study. In contrast, the Patagonian populations showed a high degree of connectivity, which could explain their recovery and high levels of current diversity. Our research emphasizes the precarious genetic status of the Uruguayan sea lion population, calling for the immediate implementation of conservation measures.

## Introduction

The spatial distribution of genetic diversity within species and the evolutionary processes involved are key features for their conservation (Moritz, 2002; Hewitt, 2004). In particular, understanding how species became subdivided into genetic units is of utmost importance because these reservoirs have resulted from evolutionary processes acting at the intraspecific level. Enhancing their genetic diversity could increase the ability of the species to survive environmental changes (Wang, 2002). In turn, one of the main objectives of conservation is to preserve the evolutionary potential of species by maintaining the diversity found in genetic units (Moritz, 1994).

Historically, human interaction with marine ecosystems has dramatically impacted marine populations, and most marine mammals have experienced severe overharvesting over the past 300 years (Busch, 1987). For example, unregulated harvesting by either deliberate or incidental bycatch led to reduced genetic diversity in Hector’s dolphin *Cephalorhynchus hectori* (Pichler and Baker, 2000); the North Atlantic right whale *Eubalaena glacialis* (Waldick *et al*., 2002); the South American endemic franciscana dolphin *Pontorporia blainvillei* (Gariboldi *et al*., 2016); and the southern right whale *Eubalaena australis* (Carroll *et al*., 2020), among others. Likewise, pinniped species have been heavily exploited worldwide, causing the decline or extinction of different populations over the last two centuries (Bonner, 1982; Gerber and Hilborn, 2001; Kovacs *et al*., 2012; Hoffman *et al*., 2016). However, the prohibition or regulation of commercial sealing had a positive effect on the recovery of these populations; in some cases, their abundance returned to pre-harvesting levels or levels compatible with available food resources and suitable breeding habitats (Franco-Trecu *et al*., 2015).

The South American sea lion, *Otaria flavescens* (Shaw, 1800) (Figure 1), was one of the most intensively hunted pinnipeds for its meat, oil and fur during the 19th and 20th centuries (Crespo et al., 2009; Godoy, 1963; Páez, 2006). It is distributed along approximately 10,000 km of the coast of South America, from Recifes das Torres (29°20’S, 49°43’W) in southern Brazil to Cape Horn in the extreme south of the Atlantic coast, and from Cape Horn to Zorritos (3°40’S, 80°34’W) in northern Peru in the Pacific Ocean (Cappozzo and Perrin, 2009). The most severely affected reproductive areas were located on the Atlantic coast, and sealing occurred mainly during the first half of the 20th century (Hamilton, 1939; Godoy, 1963; Dans *et al*., 2004). At least 47,000 sea lions were killed in the Uruguayan colonies between 1965 and 1991 (Ponce de León, 2000; Páez, 2006) and ∼60,000 sea lions at Malvinas (Falkland) Islands between 1935 and 1966 (Thompson *et al*., 2005; Baylis *et al*., 2015). In Argentina, harvest was conducted along the coasts of northern Patagonia (∼210,000 animals) and Tierra del Fuego (∼149,000 animals) between 1920 and 1960, while ∼39,000 animals were sacrificed in Santa Cruz province (south Patagonia). It is interesting to note that the current trends of the Atlantic Ocean populations are not correlated with the levels of hunting pressure they suffered in the past. Thus, the populations from Malvinas Islands, Patagonia and Tierra del Fuego show positive growth rates, at least since the 90’s (Thompson et al., 2005; Crespo et al., 2009; Grandi et al., 2015; Milano et al., 2020), while the Uruguayan colonies experience a constant decline in population size at an average annual rate of ∼-2% (Páez, 2006; Franco-Trecu *et al*., 2018).

**Figure 1.**
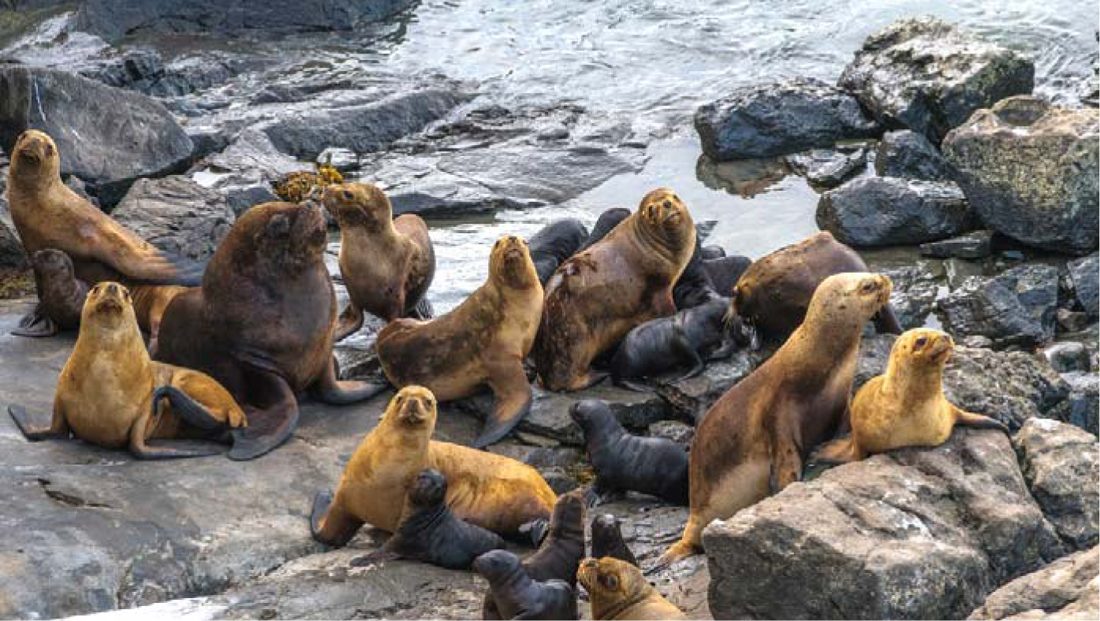
South American sea lions resting on a beach in Tierra del Fuego. The group comprises one male and several females and pups.

Currently, the species is listed as Least Concern (LC) in the IUCN Red List (Cárdenas-Alayza et al., 2016). However, this classification may be misleading because the species is partitioned into distinct genetic reservoirs or conservation units along its extensive distribution range, with particular problems requiring different solutions (Cappozzo and Perrin, 2009; Kovacs *et al*., 2012).

Many studies on population genetics and phylogeography have focused on the degree of diversity and connectivity among different colonies of South American sea lions from the Atlantic coasts (Artico et al., 2010; Feijoo et al., 2011; Hoffman et al., 2016; Oliveira et al., 2017; Peralta et al., 2021; Túnez et al., 2007, 2010). They revealed the presence of four distinct populations or conservation units (Uruguay, Patagonia, Islas Malvinas (Falklands) and Tierra del Fuego) interconnected by the colonies of Buenos Aires province (composed of males from Uruguay and northern Patagonia) and southern Santa Cruz (composed of males and females from continental Patagonia and Tierra del Fuego). Moreover, these studies provided evidence that Pleistocene glacial cycles had a strong influence on their history. Regarding the conservation status, these works found high levels of genetic diversity and structure for the Argentinean populations, using mtDNA, but low levels of genetic structure for microsatellites, possibly due to the dynamics of male migration. Feijoo *et al*. (2011) and Hoffman *et al*. (2016) failed to find evidence of bottleneck events in north-central Patagonia and Malvinas Islands. In the case of the Uruguayan population, both mtDNA and microsatellite markers detected low genetic diversity (Túnez *et al*., 2007; Oliveira *et al*., 2017). However, the decline in this population remains a mystery for population geneticists. Nevertheless, it has been attributed to long-term commercial sealing (Vaz-Ferreira *et al*., 1984), fishery interactions (De María *et al*., 2014), disruption of the social structure (Franco-Trecu *et al*., 2015), or unknown reasons (Páez, 2006).

The emergence of cutting-edge technologies such as next-generation sequencing (NGS) has allowed us to dig deeper into the genomes of different species. NGS improved our understanding of their evolutionary patterns and signatures and became a valuable tool to supplement and refine conservation efforts (Janjua *et al*., 2019). Among these technologies, restriction-site associated DNA sequencing (RAD-seq) is particularly effective for non-model organisms (Baird *et al*., 2008). RAD-seq involves reduced-representation genome sequencing (Miller *et al*., 2007; Baird *et al*., 2008) and further discovery of thousands of genome-wide SNPs (Davey and Blaxter, 2010). Thus, it emerges as a powerful tool for studying endangered populations with limited genetic diversity or where large sample sizes are difficult to obtain (Lemopoulos *et al*., 2019).

Based on the considerations mentioned above, we aimed to analyze the genetic variability and structure in some of the most exploited populations of South American sea lions from the Atlantic Ocean using the RAD-seq method. We then discussed the possible historical scenarios that may have led to the observed pattern and the implications of our results on the current conservation status of these populations after heavy harvest pressure.

## Methods

### Samples

We used a subset of 70 individuals from Peralta *et al*. (2021) to study three heavily exploited populations of *O. flavescens* in the South Atlantic Ocean (Uruguay, Patagonia, and Tierra del Fuego). After filtering RAD-seq data and assigning individuals to populations (see below), the final dataset included 32 samples from Uruguay (n=5), North-Central Patagonia (PatNC; n=14), Southern Patagonia (PatS; n=6) and Tierra del Fuego (TDF; n=7) (Table 1) (Figure 2). Samples assigned to the Uruguayan population were obtained from the locality of Puerto Quequén, Buenos Aires province, a non-reproductive colony where males from the Uruguayan and Patagonian populations rest after the reproductive season (Giardino *et al*., 2016; Peralta *et al*., 2021). However, since we cannot assign these samples to any natal breeding colony with 100% certainly, their origin was based on location and knowledge of movement from a prior tagging research (Giardino *et al*., 2016) and on the genetic results. The colonies from the Patagonian population were split into two groups (PatNC and PatS) according to the different harvesting levels they suffered (Túnez *et al*., 2008); moreover, the results from Admixture support this differentiation. Each individual was assigned to each population by estimating individual co-ancestry with ADMIXTURE v.1.3 (Alexander, Novembre and Lange, 2009). We performed 10 multiple runs for seven possible groups (K = 1 to 7), using 19,858 SNPs after pruning for Linkage disequilibrium and with only one SNP per locus. We run Pong (Behr *et al*., 2016), to merge and visualize the clustering outputs of the different Admixture runs. The SNPs were pruned for LD using Plink v1.9 (Purcell *et al*., 2007), with the filtering option--indep-pairwise 10 10 0.5. We selected the values of sliding window and r2 based on the results of PopLDdecay (Zhang *et al*., 2019) (Figure S1), a tool for linkage disequilibrium decay analysis.

**Figure 2.**
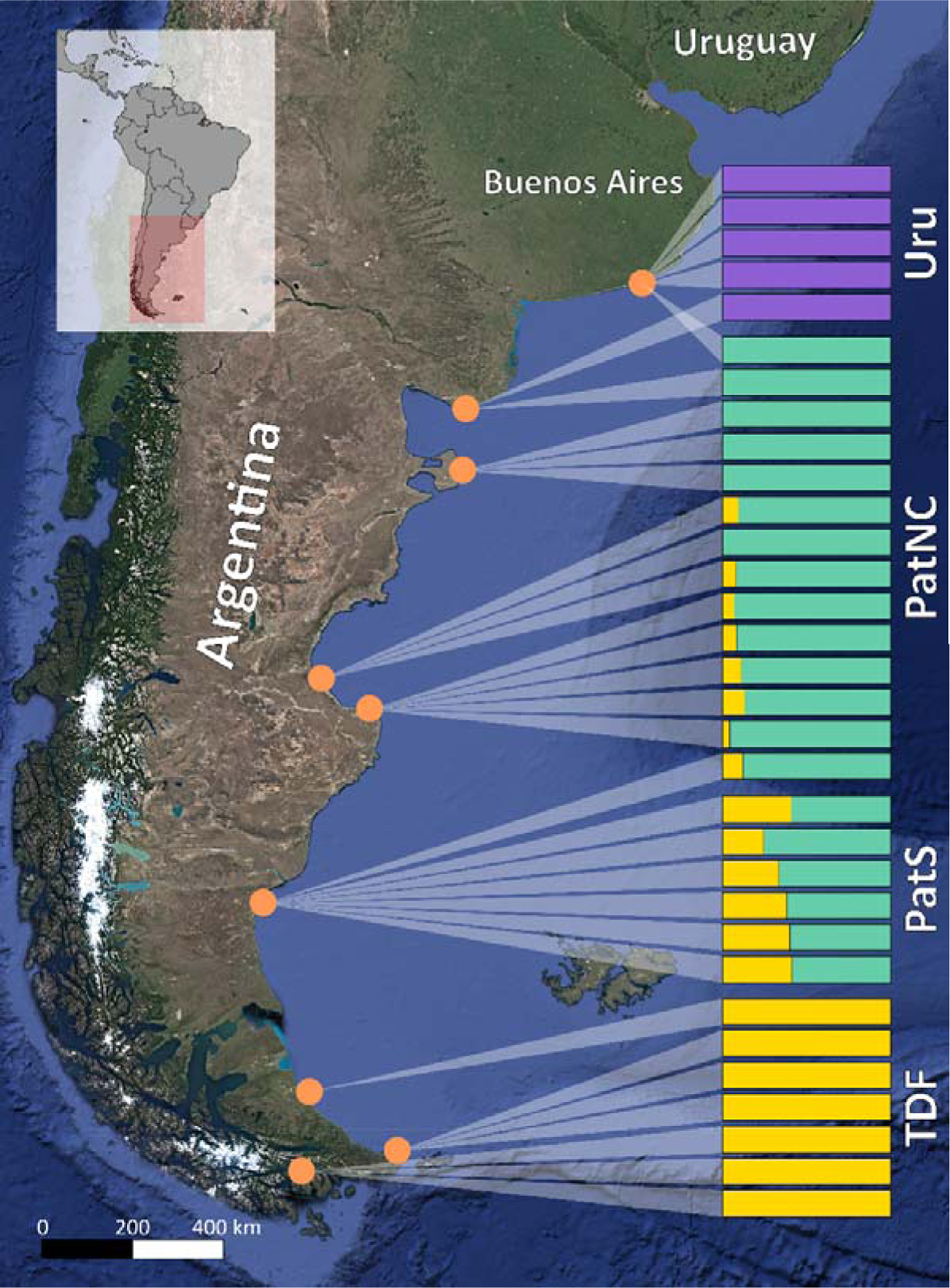
Map of the study area within the distribution range of O. flavescens in Argentina. Dots indicate sampled colonies. Shaded triangle indicates the origin of each individual and color bars represent its assignment to a particular genetic cluster based on ADMIXTURE analysis for K=3 using 19,858 LD pruned SNPs. The four studied groups of sea lions from the Atlantic Ocean are indicated on the right. The samples belonging to the Uruguayan population were collected from a non-reproductive colony in Buenos Aires province (see Methods). The Patagonian population was divided into North-Central Patagonia (PatNC) and Southern Patagonia (PatS) to better understand the species’ connectivity along the southern Atlantic coast.

**Table 1.** Summary statistics of genetic diversity in the studied *Otaria flavescens* populations.

| | N | PRIV | HO | HE | PI | $F_{IS}$ | AR | A PR |
| --- | --- | --- | --- | --- | --- | --- | --- | --- |
| URU | 5 | 44204 | 0.0581 | 0.06001 | 0.06681 | 0.01923 | 1.22654 | 0.631361 |
| PATNC | 14 | 279 | 0.11168 | 0.1248 | 0.12966 | 0.0475 | 1.54974 | 0.020559 |
| PATS | 6 | 1 | 0.11968 | 0.12218 | 0.13361 | 0.03123 | 1.40518 | 0.009917 |
| TDF | 7 | 8 | 0.09192 | 0.10835 | 0.12137 | 0.06427 | 2.11421 | 0.563509 |

Summary statistics for all sampling locations (n = 32) across 71443 biallelic SNPs. N, sample size; Priv, number of private alleles; Ho, observed heterozygosity; He, expected heterozygosity; PI, nucleotide diversity; $F_{IS}$ , inbreeding coefficient; AR, allelic richness; A PR, private allelic richness. URU: Uruguay; PATNC: Northern-Central Patagonia; PATS: Southern Patagonia; TDF: Tierra del Fuego.

### RAD sequencing

We conducted restriction site-associated DNA sequencing (RAD-seq) (Baird *et al*., 2008) following the protocol described by Roesti *et al*. (2012; 2013) and adopted by Hohenlohe *et al*. (2010). Briefly, each sample was individually subjected to restriction digestion using the Sbf1 enzyme, followed by the fusion of a specific 7-mer barcode to the restricted DNA of each individual. Two individual samples were included twice in the library for further evaluation of the genotyping error, as suggested by Mastretta-Yanes *et al*. (2015). The RAD-seq library was single-end sequenced to 150 bp in a single lane on an Illumina HiSeq 4000 at the Genomics and Cell Characterization Core Facility, Oregon University, USA.

### SNP calling and filtering

Reads quality was assessed using FastQC v0.10.1; sequences were demultiplexed using *process_radtags* in STACKS v2.53 (Catchen *et al*., 2013) and trimmed to 143 pb (after removing the 7-bp barcode and the last nucleotide). Individual reads were mapped to the California sea lion reference genome mZalCal1.pri.v2 (GCA_009762295.2) using BWA MEM v0.7.17 (Li and Durbin, 2010) with default parameters. Any unmapped reads were removed from the SAM alignment files using SAMtools v1.7 (Li, 2011). We excluded all individual samples with an average coverage depth below 10X for further procedures. SNP calling was achieved using *gstacks* in STACKS v2.53. Output files for subsequent analysis were obtained using the module *populations* in STACKS v2.53 and the parameter settings: *--max-obs-het* 0.7, *--min-mac* 6, *-R* 0.8.

### Population genomics and structure

Population genomic statistics of variability were calculated for the four groups. Private alleles, observed and expected heterozygosity (Ho and He, respectively), nucleotide diversity (π), and inbreeding coefficient ( *F*_IS_) were calculated using the *populations* module of STACKS (Catchen *et al*., 2013). Allelic richness (Ar) and private allelic richness (A pr) were calculated using a rarefaction approach implemented in ADZE v 1.0 (Szpiech *et al*., 2008). The rarefaction method calculates the expected number of alleles at a locus for a fixed sample size, generally considering the smallest sample size in a series of sampled populations (Petit *et al*., 1998) (Figure S2). Private allelic richness differs from merely counting private alleles because it allows standardizing the count of private alleles since rarefaction compensates for differences in sample size among sites. Thus, it avoids counting a larger number of private alleles at sites with larger sample sizes. (Kalinowski, 2004).

We inferred population structure by three different approaches. First, we estimated the genetic differentiation by estimating the pairwise fixation index (Fst) according to (Weir and Cockerham, 1984) with the stamppFst function from R the StAMPP package (Pembleton and Pembleton, 2015). Second, we used the *fineRADstructure* software package (Malinsky *et al*., 2018), which constructs a co-ancestry matrix of haplotypes through a Markov chain Monte Carlo (MCMC) clustering algorithm and quantifies nearest neighbor relationships between all individuals in the analysis. This approach measures recent co-ancestry by focusing on the most recent coalescent events among the haplotypes at each locus. High values, indicating a high probability of shared co-ancestry, are clustered along the diagonal, and these individuals are clustered into populations in the accompanying tree. We used a 100,000 burn-in followed by 100,000 MCMC steps, sampling every 1000 steps, and constructed the tree with 1000 hill-climbing iterations. Only autosomal SNPs (n= 69,599) were used in this analysis. We visualized and plotted the results using R scripts fineRADstructurePlot.R and FinestructureLibrary.R (available at http://cichlid.gurdon.cam.ac.uk/fineRADstructure.html).

Last, we performed a principal component analysis (PCA) using the R package adegenet (Jombart and Ahmed, 2011) with the glPCA function. We only used autosomal SNPs (n=69,599) and conducted two different analyses to infer population structure; one included the whole data set and the other excluded data from Uruguay.

## Results

### SNP discovery

After filtering for quality and coverage depth, we obtained an average of 3.2 million reads per sample. A total of 138,784 widely shared loci (present in at least 80% of samples) were obtained with an average depth of 21.2 X ± 6.76. Further filtering using *population* module with --min-mac 6 retained 71,443 SNPs belonging to 118,171 shared loci, which were used for subsequent analysis. The estimated SNP error rate for this dataset in two replicated sample pairs was 0.0123 ± 0.0038.

### Population genomics and structure

The genetic ancestry inferred by ADMIXTURE was used to assign each individual to its original population, as shown in Figure 2. Individuals from the non-reproductive colony of Buenos Aires province were reassigned to Uruguay or Northern Patagonia. Both, the cross-validation error method implemented in ADMIXTURE and the Pong’s Jaccard similarity index indicate K=2 and K=3 as the most plausible scenarios for explaining the genetic variability (Figure S3, Figure S4), which is consistent with a hierarchical population structure. Finally, we assigned individuals to four groups representative of the studied populations, as explained in Methodology; i.e., Uruguay (Uru), North-Central Patagonia (PatNC), Southern Patagonia (PatS) and Tierra del Fuego (TDF).

Measures of Expected and Observed heterozygosity (He and Ho, respectively) and nucleotide diversity (π) were similar in PatNC, PatS and TDF (ranging between 0.09-0.12 for Ho; 0.10-0.12 for He; and 0.12-0.13 for π, respectively) but lower for Uru (Ho=0.06, He=0.06 and π=0.06). In all cases, the ratio Ho/He was about 1. *F*_IS_ coefficients differed among groups, ranging between 0.019 and 0.064. Uru showed the lowest *F*_IS_ coefficient and TDF the highest one. Uru had 44,204 private alleles, and the other groups had between 1 and 279 unique alleles. Allelic richness was highest for TDF (Ar = 2.11, stderr = 0.31) and lowest for Uru (Ar = 1.22, stderr = 0.19), while the two Patagonian groups exhibited intermediate values (Ar= 1.405 and stderr= 0.33 for PatNC; Ar= 1.55 and stderr= 0.302 for PatS). Private allelic richness was highest for Uru (A pr = 0.63, stderr = 0.23) and lowest for the groups of Patagonia (A pr = 0.01 and stderr = 0.004 for PatNC; A pr = 0.02 and stderr = 0.01 for PatS). TDF also showed high private allelic richness (A pr = 0.56, stderr = 0.1829) (Table 1).

Pairwise comparisons between Uruguay and all other groups yielded the highest *F*_ST_ values (i.e., 0.84 versus PatNC, 0.86 versus PatS, and 0.87 versus TDF). The FST values of the comparisons between the remaining groups were lower than 0.078 (Table 2). All the comparisons presented significant differences (P < 0.001).

**Table 2.** Pairwise Fixation index *F*_ST_ among the four assigned groups of the studied *Otaria flavescens* populations. All the comparisons presented significant differences (P < 0.001).

|  | Uru | PatNC | PatS | TDF |
| --- | --- | --- | --- | --- |
| Uru | - | 0.84821 | 0.85894 | 0.86979 |
| PatNC |  | - | 0.013204 | 0.077947 |
| PatS |  |  | - | 0.033389 |
| TDF |  |  |  | - |

Population acronyms as in Table 1.

Results from *FineRADstructure* indicated two main genetic groups, one comprising individuals from Uruguay and the other individuals from the Argentinean colonies. Within the Argentinean group, individuals of PatNC were distinguished from those of TDF, albeit at a lower level of co-ancestry (Figure 3.A). However, the exclusion of Uruguayan individuals from the analysis allowed a better resolution among the populations of Argentina. This new matrix led to the detection of a higher level of co-ancestry among the individuals of TDF, and to a lower extent, among individuals of Patagonia. In addition, the latter analysis indicated that individuals from PatS were equally related to those from PatNC and TDF (Figure 4.A).

**Figure 3.**
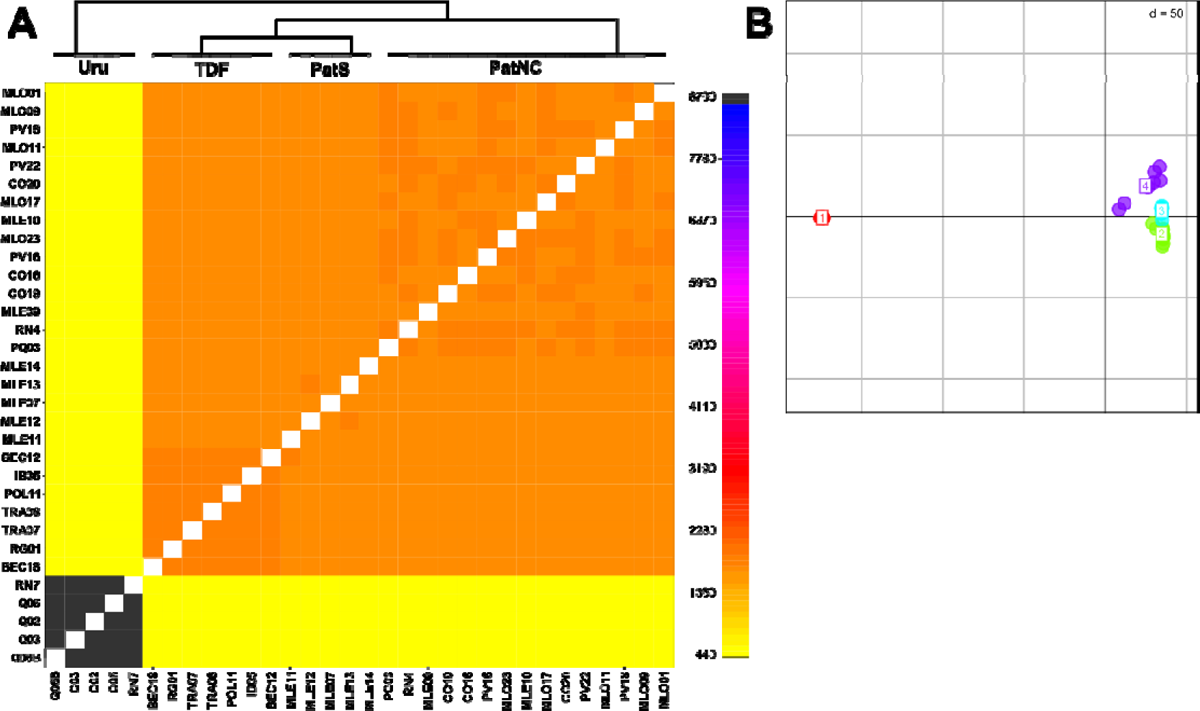
fineRADstructure and Principal Component Analysis (PCA) for the full dataset. A. The co-ancestry matrix indicates pairwise co-ancestry between individuals. B. First two PCA components; PC1 explains 73% of the total variation. Red dots correspond to Uru; green dots correspond to PatNC; light blue dots correspond to PatS; violet dots correspond to TDF.

**Figure 4.**
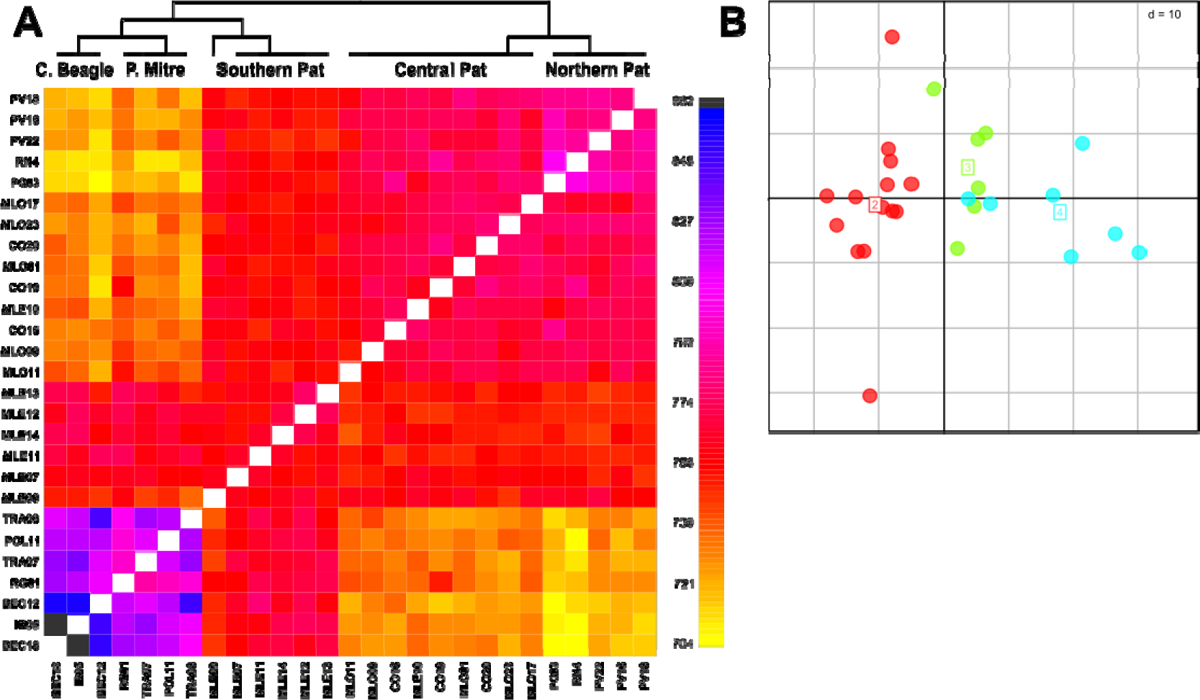
fineRADstructure and Principal component analysis (PCA) for the populations of Argentina. A. The co-ancestry matrix indicates pairwise co-ancestry between individuals. B. First two PCA components; PC1 and PC2 explain 7.71% and 4.28% of the total variation, respectively. Red dots correspond to PatNC; green dots correspond to PatS; light blue dots correspond to TDF.

The PCA of SNPs from autosomal chromosomes showed two divergent groups, one composed of individuals from Uru and the other from Patagonia and TDF (Figure 3.B). Once more, the structure of the Argentinean colonies was better resolved after Uru was removed from the analysis, with PatS showing an intermediate position between PatNC and TDF (Figure 4.B).

## Discussion

To our knowledge, this is the first study on the population genomics of the South American sea lion using RAD-seq and the first time that this methodology has been applied to study a South American pinniped. Here, we took advantage of the high-resolution power of this technique to improve our understanding of the history and connectivity among three overhunted populations, and their current conservation status.

### Population assignment of individuals

The identification of the population source of each individual was performed using ADMIXTURE. In most pinnipeds, it is unlikely that all individuals from a colony belong to the same population due to the high philopatry of females and the seasonal migratory dynamics of males (Perrin and Mazalov, 2000; Giardino *et al*., 2016). Thus, this step allows us to avoid misleading results caused by grouping individuals from different genetic clusters into a single one, a common issue to consider in highly vagile species. Moreover, the analysis presented a first visualization of the hierarchical population structure of this species in the Atlantic Ocean. A primary genetic differentiation between the Argentinian and the Uruguayan populations (with K=2), but also the differentiation within Argentina at a lower hierarchical level (only evident with K=3) (Figure 2).

### Population diversity and differentiation

Genetic diversity is an important aspect of population dynamics, as it is directly related to the evolutionary potential of a population and the deleterious effects of inbreeding (Hughes *et al*., 2008). In this regard, heterozygosity and allelic richness are the most commonly used measures to test for loss of genetic diversity (Greenbaum *et al*., 2014). Loss of heterozygosity is a good indicator of a decrease in the average fitness of individuals (Reed and Frankham, 2003; Szulkin et al., 2010). In comparison, the loss of allelic richness indicates the reduction in populations’ potential capacity to adapt to future environmental changes since this diversity is the raw material for evolution by natural selection (Fisher, 1958). Moreover, allelic richness is frequently regarded as the most useful neutral measure for monitoring changes in population genetic variation because it is susceptible to population changes (Phair *et al*., 2020).

Our analyses of diversity indicated low levels of heterozygosity, nucleotide diversity and allelic richness for the Uruguayan population. These results rated it as the least variable in the Atlantic Ocean, in accordance to previous works reporting a constant decline in population size and low genetic variability (Páez, 2006; Túnez *et al*., 2007; Oliveira *et al*., 2017; Franco-Trecu *et al*., 2018). North-Central and Southern Patagonian populations showed moderate to high diversity values, while Tierra del Fuego presented a combination of high allelic richness and moderate/low heterozygosity.

On the one hand, the low variability found in Uruguay may result from low reproductive success, as suggested by Franco-Trecu *et al*. (2015) and a weak genetic load. However, on the other hand, the Tierra del Fuego population, showed the highest allelic richness and private allele richness, but with similar Pi and Ho values to those in Patagonia. According to our last findings, the most likely explanation for the high genetic diversity of TDF could be related to its recent history and to the secondary contact between the lineages of the Atlantic and Pacific Oceans after the end of the last glacial period (Peralta *et al*., 2021). However, the mentioned difference between allelic richness and heterozygosity observed in this population may be due to the massive overhunting suffered in the last century (Túnez *et al*., 2008) and subsequent recovery.

With respect to population structure, RAD-seq analysis revealed that the Uruguayan population was isolated from the other studied populations. Pairwise *F*_ST_ values showed a high genetic differentiation between the Uruguayan and the Argentinean populations, even considering PatNC, which has colonies located only 600 km apart from Uruguay. In comparison, Patagonian groups and TDF exhibited a high degree of connectivity.

Following the results presented above, *fineRADstructure* analysis provided evidence of complete isolation for Uru. The high degree of co-ancestry within this population is consistent with the low diversity indicated by the analyses of heterozygosity, all elicrichness and nucleotide diversity. The level of differentiation measured by the co-ancestry matrix suggests the absence of gene flow between the Uruguayan and the Argentinean populations. On the contrary, the PatNC and TDF clusters found in the second matrix seems to have a high degree of gene flow mediated by the colonies in PatS. It is worthwhile to note the extremely high resolution capacity of *fineRADstructure* in the second matrix, as it permitted to cluster individuals according to their precise geographical origin and even to differentiate genetically between the northern and southern coasts of TDF. Thus, the accompanying tree discriminated between samples from Canal Beagle and Península Mitre in TDF and between samples from the north and center of Patagonia.

PCA results agree with those of *fineRADstructure* showing a strong genetic differentiation for the Uruguayan population but a closer genetic relationship among Patagonian groups and Tierra del Fuego. In summary, we found a high genetic differentiation between Argentina and Uruguay and a subtle differentiation within Argentina, where PatS would share intermediate co-ancestry with PatNC and TDF. Túnez *et al*. (2007) and Peralta *et al*. (2021) suggested that the latter pattern may be explained by the Stepping Stone model, which states that most gene flow occurs between closer colonies.

Previous studies reported a low genetic variability for the Uruguayan population, but overlooked its extreme isolation from the other Atlantic populations. Túnez *et al*. (2007) found a unique cyt B haplotype in colonies from Uruguay against six haplotypes found across the species distribution. Oliveira *et al*. (2017) observed that this population had moderate to high levels of diversity for the mtDNA control region; however, it showed a lower number of alleles and a remarkable difference between Ho and He for microsatellites when compared to the Argentinean populations. Regarding genetic structure, this work would have shown a glimpse of what we found here. Even though they only found significant genetic differences between the Uruguayan and Argentinean populations using the mtDNA marker, their microsatellites showed genetic divergence for the Uruguayan population within the Atlantic Ocean.

The indices of genetic diversity calculated by RAD-seq in the present paper do not seem comparable with those obtained with microsatellites in previous works. Some studies have shown weak positive correlations between these techniques (Väli *et al*., 2008; Glover *et al*., 2010; Ciani *et al*., 2013), but results strongly depend on the number of SNPs. Studies using more than 3000 SNPs reported that these markers are better for inferring genetic diversity (Väli *et al*., 2008; Glover *et al*., 2010; Gärke *et al*., 2012; Fernández *et al*., 2013; Ozerov *et al*., 2013). Another advantage of using a large data set of SNPs over microsatellites is their power to resolve the genetic structure. Jeffries *et al*. (2016), who used microsatellites with a large set of samples versus SNPs from RAD-seq using 17% of these samples, found that the latter had a better performance for recovering finer population structure. In our case, despite the small sample size in the present study, the degree of definition afforded by our large marker dataset (70,000 SNPs) makes our results highly reliable. Therefore, the approach used here provides a more accurate perspective on the conservation status and genetic structure of Atlantic sea lion populations, solving previous unknowns.

### Possible explanations for the genetic signature

The phylogeography of *O. flavescens* has intrigued many scientists since the first publication by Szapkievich *et al*. (1999) using allozymes. Inthissense, our results raisean interesting question: What would have been the reasons that caused the Uruguayan population to be isolated from Patagonia? Peralta *et al*. (2021) using mtDNA, suggested that Uru population might be the older and most diverse population in the Atlantic Ocean, as well as the source of origin for the current southern populations after the end of glaciations at approximately 150,000 years ago.

However, our new results indicate that at some point in its history it would have become completely isolated from the other populations, followed by a drop in diversity levels. The contrasting scenarios obtained with the two different molecular markers led us to assume that its current situation is the result of events occurring in the Late Pleistocene (e.g., the Last Glacial Maximum) and thousands of years of separation causing complete lack of gene flow.

Interestingly, the separation of Uruguay occurred at the same time that the Patagonian and Tierra del Fuego populations were becoming very diverse (mainly TDF) and highly interconnected. Such promising variability levels and positive population trends could be due to their proximity and the high level of gene flow in the area, as suggested by fineRADstructure. Altogether, these situations may explain the differences in diversity and structure between the southern Atlantic populations and that in Uruguay.

Another important aspect to consider is the extreme overhunting of the Atlantic populations during the last two centuries. Based on records for the period of greatest commercial harvest (∼1917-1991) and population censuses carried out between 1950 and 1995, the overexploitation of *O. flavescens* in Uruguay, northern Patagonia and Tierra del Fuego led to a population decline of more than 50%. Tierra del Fuego and Northern Patagonia were the most affected areas, (95% and 70%, respectively). However, current population trends do not correlate with the levels of hunting pressure. Patagonia and Tierra del Fuego show positive growth rates (Crespo *et al*., 2009; Grandi *et al*., 2015; Milano *et al*., 2020), while the Uruguayan colonies experience a constant decline at an average annual rate of ∼-2% (Páez, 2006; Franco-Trecu *et al*., 2018). We postulate that the high level of connectivity in the area, together with the dissimilar harvest levels across colonies of each population (Crespo and Pedraza, 1991; Sielfeld, 1999) might have counteracted the impact of the high exploitation rates on the southern populations, while this was impossible in the genetically isolated Uruguayan population. Indeed, Oliveira *et al*. (2017) suggested that the elimination of all breeding colonies of Buenos Aires during the sealing of XVIII could explain the genetic difference they found with mitochondrial markers.

Our hypothesis is supported by evidence indicating that high levels of connectivity have a compensatory effect on declining populations (Lowe and Allendorf, 2010), with the response being even stronger when individuals come from a more genetically diverse population (Lansink *et al*., 2022). We therefore propose that like for other vagile species, the recovery of depleted populations in *O. flavescens* would be closely tied to their degree of connectivity with neighboring populations. In line with this, our study highlights the key role that Patagonia and Tierra del Fuego colonies play as potential sources of genetic diversity for the recovery of depleted sea lion populations through immigration.

## Conclusions

This study compares the genetic connectivity and conservation status among three of the most extensively harvested populations of *O. flavescens*. We found moderate to high levels of genetic diversity and soft genetic structure between the Patagonian and Tierra del Fuego populations. Moreover, our results disclosed, for the first time, the precarious condition of the Uruguayan population due to its remarkable genetic isolation and depletion, and we postulate that it would have been isolated from Patagonia since the Late Pleistocene. Indeed, a comparison with the southern populations led us to hypothesize that the lack of gene flow may help explain its failure to recover. However, we cannot rule out the possible influence of climatic, ecological and/or anthropological factors on its current situation. Therefore, further studies combining data on these factors with genetic information and the demographic changes that this population has undergone in the recent past may help elucidate this issue.

Finally, results emphasize the importance of connectivity in the recovery of depleted populations, and like in previous studies, we call for the urgent implementation of conservation measures for the Uruguayan population.

## Acknowledgments and funding information

We are grateful to the park rangers of the National Parks and Protected Areas for their cooperation and hospitality; to the government agencies and their staff for issuing the necessary permits (Administración de Parques Nacionales, Secretaría de Medio Ambiente y Desarrollo Sustentable de Río Negro, Secretaría de Ambiente de Santa Cruz and Secretaría de Ambiente, Desarrollo Sostenible y Cambio Climático de Tierra del Fuego); to the staff of the NGO Fundación Cadace for their support in Caleta Olivia; to Adolfo Imbert, Dario Urruti and Agustín “Chepan” Szczepañsk from Centro Hípico del Fin del Mundo, Ushuaia, for guiding us in Península Mitre; to Lida Pimer for helping with local permits and logistics; to Proyecto IMMA for helping us with samples from Beagle Channel; and to Centro Austral de Investigaciones Científicas. This work was supported by fellowships from CONICET and DGAPA-UNAM, and grants from the National Geographic Society (10000-16 and EC-52880R-18) and the Universidad Nacional de Luján (Finalidad 3.5 año 2017). The authors declare that they have no conflict of interest. Last, we also want to thank to the anonymous reviewer for their helpful comments.

## Data accessibility

Sequence data will be deposited in Gen Bank upon publication.

